# *Escherichia coli* Cryptic Prophages Sense Nutrients to Control Persister Cell Resuscitation

**DOI:** 10.1101/2021.04.10.439273

**Authors:** Sooyeon Song, Jun-Seob Kim, Ryota Yamasaki, Sejong Oh, Michael Benedik, Thomas K. Wood

**Author notes:** For correspondence. (+)1.

## Abstract

We determined previously that some cryptic prophages are not genomic junk but instead enable cells to combat myriad stresses as part of an active stress response. However, how these phage fossils affect the extreme stress response of dormancy; i.e., how cryptic prophages affect persister cell formation and resuscitation, has not been fully explored. Persister cells form as a result of stresses such as starvation, antibiotics, and oxidative conditions, and resuscitation of these persister cells likely causes recurring infections such as those associated with tuberculosis, cystic fibrosis, and Lyme disease. Unlike for the active stress response, here we find that deletion of each of the nine *Escherichia coli* cryptic prophages has no effect on persister cell formation. Strikingly, elimination of each cryptic prophage results in an increase in persister cell resuscitation with a dramatic increase in resuscitation upon deleting all nine prophages. This increased resuscitation includes eliminating the need for a carbon source and is due to activation of the phosphate import system as a result of inactivating transcriptional regulator AlpA of the CP4-57 cryptic prophage, since we found Δ*alpA* increases persister resuscitation, and AlpA represses phosphate regulator PhoR. Therefore, we report a novel cellular stress mechanism controlled by cryptic prophages: regulation of phosphate uptake which controls the exit of the cell from dormancy and prevents premature resuscitation in the absence of nutrients.

## INTRODUCTION

We have sought to understand the role of trapped lysogens that no longer form active phage particles, i.e., cryptic prophages, since bacterial genomes may have up to 20% phage DNA^1^ and much of this DNA is stable. Rather than merely being extraneous DNA, we found that the nine cryptic prophages of *E. coli* K-12 increase resistance to sub-lethal concentrations of quinolone and β-lactam antibiotics as well as protect the cell from osmotic, oxidative, and acid stresses^2^. Hence, the genome of a parasite has become interwoven into the bacterial genome and serves beneficial roles related to stress resistance^2^. Since the extreme response of the cell population to stress is for some cells to become dormant (i.e., persistent)^3^ and cryptic prophages are utilized for an active stress response^2^, we reasoned that cryptic prophages may play a role in persistence; i.e., cryptic prophages may be involved in the extreme stress response.

Persister cells are phenotypic variants that arise without genetic change as a result of myriad stresses such as nutrient, antibiotic, and oxidative stress^4, 5^. Most cells in the stressed population mount an active response but the small subpopulation of persisters survive stresses by sleeping through the insults^6^. Since nearly all cells starve, the persister state is probably a universal resting state^7^. Critically, persister cells are likely the cause of many recurring infections^8^; therefore, understanding how they resuscitate is vital.

We have determined that persister cells can form as a result of translation inhibition. Specifically, by inhibiting transcription via sub-lethal concentrations of rifampicin, by inhibiting translation via sub-lethal concentrations tetracycline, or by interrupting translation by ceasing ATP production via carbonyl cyanide *m*-chlorophenyl hydrazone, we converted nearly all of the exponentially-growing *E. coli* cells into persister cells^9^. To reduce protein production during stress, there is consensus^10-12^ for a role of the alarmone guanosine pentaphosphate/tetraphosphate (henceforth ppGpp) for forming persisters^13^; for example, by reducing transcription of ribosomal operons^14^. We have proposed the ppGpp ribosome dimerization persister (PRDP) model in which ppGpp generates persister cells directly by inactivating ribosomes through the conserved ribosome hibernation factors RMF, HpF, and RaiA which convert active 70S ribosomes into inactive 70S and 100S ribosomes^15-17^. In support of this model, we found that persister cells contain a large fraction of 100S ribosomes, that inactivation of RMF, HpF, and RaiA reduces persistence and increases single-cell persister resuscitation and that single-cell persister resuscitation does not depend on ppGpp^15^. This model does not rely on toxin/antitoxin systems for persister cell formation as their direct link to persistence is tenuous^18^.

We have found persister cells resuscitate in a heterogeneous manner as they recognize external nutrients; the rate of resuscitation depends on the number of active ribosomes, and the growth rate of the resuscitated cells is exponential like the wild-type^19^. Later studies have verified this heterogeneous nature of persister cell resuscitation^20, 21^ and subsequent exponential growth of resuscitating cells^22^; together, these later studies corroborate our method of forming persister cells. By screening 10,000 compounds for persister cell resuscitation, we determined that ribosomes are activated during persister cell resuscitation by pseudouridine synthase RluD that modifies 23S subunits^17^. Resuscitation is initiated by recognizing external nutrients through receptors for chemotaxis (for amino acids) and phosphotransferase membrane proteins (for glucose)^3^. The external nutrient signals are propagated to the cytosol by reducing concentrations of the secondary messenger cAMP which leads to the rescue of stalled ribosomes and to the dissociation of inactive dimerized 100S ribosomes^3^. The resuscitating cells also initiate chemotaxis toward fresh nutrients since nutrient depletion triggered persistence^3^.

Here we explore the role of cryptic prophages in persister cell formation and resuscitation. To link cryptic prophages to stress resistance, we utilized our Δ9 strain in which we precisely deleted all nine cryptic prophage elements (166 kb) of *E. coli* along with the set of single strains with each of the nine cryptic prophages deleted^2^. We find that cryptic prophages do not play a role in persister cell *formation* but instead reduce persister cell *resuscitation* and prevent resuscitation until a carbon source is present. By employing a whole transcriptome study to explore the impact of cryptic prophages on resuscitating cells, we found that the cryptic prophages reduce resuscitation by repressing phosphate transport. Moreover, we determined this phosphate transport system is repressed in part via the CP4-57 cryptic prophage regulator AlpA; specifically, AlpA represses *phoR*, which leads to activation of *phoB* and *pstB*. Hence, we discovered a novel physiological role for cryptic prophages, regulation of persister cell resuscitation, and determined that the mechanism is through their regulation of phosphate sensing.

## RESULTS

### Cryptic prophages do not affect persister cell formation

To determine whether cryptic prophages affect persister cell formation, we converted the wild-type, the Δ9 strain with all nine cryptic prophages deleted^2^, and each single cryptic prophage mutant^2^ to persister cells using a rifampicin pretreatment^9^ and enumerated them. This method of generating persister cells has been validated eight ways^19^ and utilized by at least 15 independent labs to date^9, 20, 23-35^; the approach has the benefit of increasing the number of persister cells by 10,000-fold^9^ which enables single-cell studies^3, 5, 15, 19^. We found the presence of the cryptic prophages had no significant impact on the number of persister cells formed with ampicillin (**Fig. 1**). Corroborating this result, ATP levels for persister cells were similar for both the wild-type and the Δ9 strain (**Table S1**); previously, we demonstrated reducing ATP levels increases persistence by 10,000-fold with ampicillin^9^. Hence, although cryptic prophages dramatically increase resistance to stresses^2^, they do not change the number of the cells that become persisters. These results are similar to those of Harms et al.^36^ who utilized our Δ9 strain^2^ and found little impact on the number of persister cells after treating with ciprofloxacin.

**Fig. 1.**
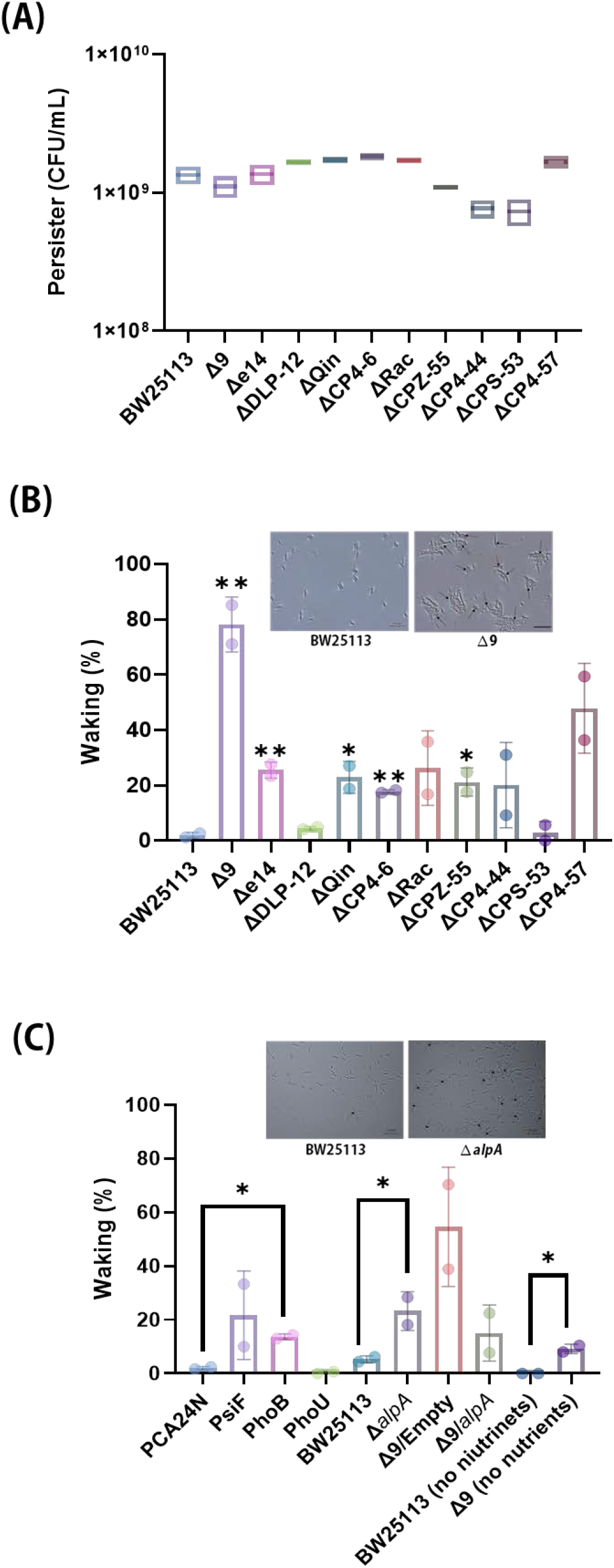
Cryptic prophages have no effect on persister cell formation but reduce single-cell persister resuscitation by repressing phosphate sensing. **(A)** Cryptic prophages do not affect persister cell formation. Persister cell formation (CFU/mL) was determined by cell counts on LB plates after 1 day. These results are the average of two independent cultures and error bars indicate standard deviations. **(B)** Cryptic prophages reduce persister cell resuscitation. Single-cell persister resuscitation after 4 h at 37°C with 0.4 wt% glucose as determined using light microscopy for each of the single cryptic prophage deletions as well as for the Δ9 strain. Representative results from two independent cultures are shown, tabulated cell numbers are shown in **Table S2**, and representative images are shown in **Fig S1**. Representative images for BW25113 and Δ9 resuscitating shown in the inset. **(C)** PhoB increases single-cell persister resuscitation. Single-cell persister resuscitation after 6 h for BW25113/pCA24N vs. BW25113/pCA24N_*psiF* (“PsiF”), BW25113/pCA24N vs. BW25113/pCA24N_*phoB* (“PhoB”), and BW25113/pCA24N vs. BW25113/pCA24N_*phoU* (“PhoU”), after 4 h for BW25113 vs. BW25113 Δ*alpA*, after 3 h for Δ9/pCA24N vs. Δ9/pCA24N_*alpA*, and after 3 h for BW25113 (no nutrients) vs. BW25113 Δ9 (no nutrients); the different resuscitation times were used to distinguish more clearly the differences in resuscitation. Cells were resuscitated at 37°C with 0.4 wt% glucose (except for the “no nutrient” group) as determined using light microscopy. Representative results from two independent cultures are shown, tabulated cell numbers are shown in **Table S3**, and representative images are shown in **Fig. S4**. Representative images for BW25113 and Δ*alpA* resuscitating shown as the inset. Student’s T-tests were used to compare pairs (* indicates a p value < 0.05 and ** indicates a p value < 0.01).

### Cryptic prophages reduce persister resuscitation

To investigate further the role of cryptic prophages in persistence, we quantified single cell resuscitation for the Δ9 strain in M9 glucose medium. In stark contrast to the lack of impact of the cryptic prophages on persister cell *formation* (**Fig. 1A**), deleting the nine cryptic prophages dramatically increased persister cell *resuscitation* (44-fold, **Fig. 1B, Fig. S1, Table S2**). Remarkably, nearly all of the persister cells that lack cryptic prophage (80%) resuscitated within 4 h compared to 2% of the wild-type strain. Similar results were obtained by treating only with ampicillin to form persister cells (i.e., foregoing rifampicin pre-treatment, **Fig. S2**) in that 66% of the single cells resuscitated in the absence of the cryptic prophages (a 34-fold increase). Furthermore, we corroborated the results with the Δ9 strain (166 kb deleted) by testing for single-cell resuscitation of *E. coli* MDS66^37^ with has 19% of the chromosome deleted (891 kb) including the cryptic prophages deleted and found this strain also had a 17-fold higher resuscitation than the wild-type (**Fig. S3, Table S2**).

We next investigated the impact of each cryptic prophage by testing each of our single complete prophage deletions^2^ and found deletion of each cryptic prophage increased the frequency of waking by 2.3 to 25-fold (**Fig. 1B, Fig. S1, Table S2**); the frequency of resuscitation was greatest upon deleting the genes of cryptic phage CP4-57 (25-fold). Hence, each cryptic prophage reduces the frequency of persister cell resuscitation although the contribution of each varies.

### Cryptic prophages repress resuscitation by reducing phosphate sensing

To elucidate the mechanism by which the cryptic prophages reduce persister cell resuscitation, we performed a whole-transcriptome analysis of resuscitating persister cells for Δ9 and compared to the wild-type under the same conditions (**Table 1**). Remarkably, we found that five phosphate-sensing genes, *pstSCB, phoR*, and *phoB*, were induced in the absence of the cryptic prophages. PhoR-PhoB comprise a two-component regulatory system that activate the phosphate-specific transport (Pst) system that includes the outer membrane ATP-binding cassette proteins PstSCAB which act as an inorganic phosphate sensor^38^. Therefore, the whole-transcriptome data suggest deletion of the cryptic prophages increases persister resuscitation by increasing phosphate sensing.

**Table 1.** Most induced genes during persister cell resuscitation for BW25113 Δ9 vs. BW25113. Persister cells were formed using the rifampicin/ampicillin method, and persister cells were resuscitated by adding M9 glucose (0.4 wt%) for 10 min. Two independent cultures were used for each strain.

| Gene | Description | Fold Change for $\Delta 9$ vs. BW25113 |
| --- | --- | --- |
| *ssrA* | transfer-messenger RNA | 8.1 |
| *envY* | DNA-binding transcriptional regulator | 6.3 |
| *rrf* | 5S ribosomal RNA | 4.4 |
| *cysI* | assimilatory sulfite reductase hemoprotein subunit | 3.9 |
| *eutA* | ethanolamine ammonia-lyase reactivating factor | 3.7 |
| *leuB* | 3-isopropylmalate dehydrogenase | 3.4 |
| *pstB* | phosphate ABC transporter ATP-binding protein | 3.4 |
| *pstC* | phosphate ABC transporter permease | 2.9 |
| *pstS* | phosphate ABC transporter substrate-binding protein | 2.9 |
| *phoR* | phosphate response regulator (sensor kinase) | 2.7 |
| *phoB* | phosphate transcriptional regulator | 3.0 |
| *mocA* | molybdenum cofactor cytidyltransferase | 3.3 |
| *cysH* | phosphoadenosine phosphosulfate reductase | 3.2 |
| *flgG* | flagellar basal-body rod protein | 3.0 |
| *nfrA* | bacteriophage adsorption protein | 3.0 |
| *leuC* | 3-isopropylmalate dehydratase large subunit | 2.9 |
| *nanM* | *N*-acetylneuraminate epimerase | 2.9 |
| *mcrB* | 5-methylcytosine-specific restriction endonuclease subunit | 2.9 |
| *paoD* | molybdenum cofactor insertion chaperone | 2.8 |

### PhoB increases persister cell resuscitation

Since the phosphate transport locus *pstSCAB* is activated by the response regulator PhoB, which in turn, is activated by the kinase activity of PhoR^39^ (**Fig. 2**), we hypothesized that the cryptic prophages contain at least one negative regulator that represses either *phoR* or *phoB*. Hence, activation of PhoR or PhoB should increase persister resuscitation.

**Fig. 2.**
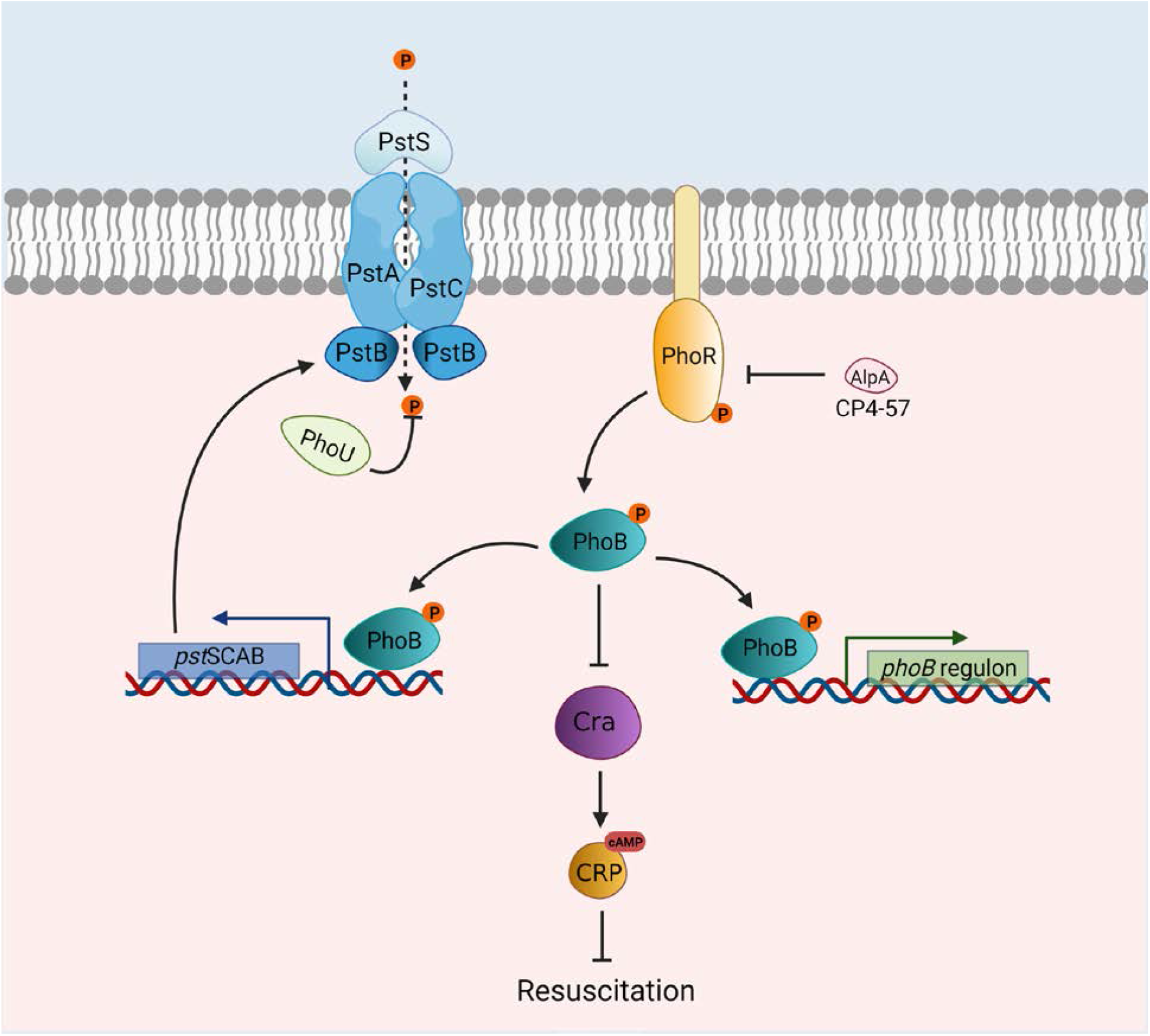
Schematic of the impact of phosphate sensing and cryptic prophage AlpA on persister cell resuscitation. AlpA (from cryptic prophage CP4-57) represses *phoR* that encodes the phosphate-dependent PhoR/PhoB two-component signal transduction system. PhoR/PhoB sense external phosphate and are active when concentrations of external phosphate are low. PhoR phosphorylates PhoB which induces the *pstSCAB* operon and facilitates phosphate uptake. PhoB increases persister cell resuscitation by reducing cAMP by silencing *cra* transcription. ⓟ indicates phosphate, → indicates induction, and ⊥ indicates repression.

To test this hypothesis, we produced PhoB from pCA24N-*phoB*; we utilized phosphate replete conditions (i.e., using M9 buffer) so that if there was an effect it would be under conditions where the phosphate transport system would normally be repressed. Under these conditions, we found that persister resuscitation was increased eight-fold by producing PhoB compared to the empty plasmid (**Fig. 1C, Fig. S4, Table S3)**. Furthermore, production of PhoU, the negative regulator of PhoR, via pCA24N-*phoU*, reduced resuscitation by four fold (**Fig. 1C, Fig. S4, Table S3**), as expected. Note we previously reported that phosphate starvation protein PsiF, which is positively regulated by PhoB, increases persistence six-fold when produced from pCA24N-*psiF*^*3*^. Hence, activation of the phosphate sensing system increases resuscitation through PhoB and PsiF, and negative regulator PhoU reduces resuscitation.

### AlpA from cryptic prophage CP4-57 represses phosphate sensing

To investigate how the cryptic prophages reduce resuscitation through phosphate sensing, we evaluated the impact of deleting three DNA putative regulators, *yfjR, alpA*, and *yfjP*, from cryptic prophage CP4-57, since this cryptic prophage had the largest impact on persister cell resuscitation. We reasoned these regulators could possibly directly or indirectly influence phosphate import. On agar plates, the *alpA* mutant woke faster than *yfjR* and *yfjP* as evidenced by consistently larger colonies; hence, we focused on the *alpA* mutant. Using single cell microscopy and six independent cultures, we found the *alpA* deletion consistently increased persister cell resuscitation by 4.4-fold (**Fig. 1C, Fig. S4, Table S3**); hence, AlpA of cryptic prophage CP4-57 reduces persister resuscitation. Moreover, production of AlpA from pCA24N complemented the *alpA* deletion by reducing persister resuscitation 4.1-fold (**Fig. 1C, Fig S4, Table S3**). Strikingly, rather than being a metabolic burden, production of AlpA *increased* the specific growth 4-fold in rich medium; hence, AlpA has a large impact on metabolism.

To explore how AlpA influences phosphate import, we used qRT-PCR to investigate AlpA activation of *phoR, phoB* and *pstB* in the absence of the nine cryptic prophages. Compared to the empty plasmid, production of AlpA via pCA24N-*alpA* in the *Δ*9 strain repressed *phoB, phoR* and *pstB* (**Table S4**); hence, AlpA likely binds to the *phoB* promoter resulting in the repression of the downstream phosphate sensing proteins PhoR and PstB (**Fig. 2**).

### Δ9 persisters resuscitate without a carbon source

Given the cryptic prophages repress resuscitation by reducing phosphate sensing, we reasoned that perhaps the role of the phage fossils was to prevent persister cell resuscitation in the absence of a carbon source. Therefore, we tested for persister cell resuscitation in the absence of glucose but in the presence of phosphate (same as M9 medium) and found the Δ9 strain resuscitates without glucose (9 ± 2%, **Fig. 1C, Fig. S4, Table S3**) whereas the wild-type strain cannot resuscitate without glucose, as we found previously for the absence of LB medium^19^ and alanine^3^. Note this increased waking by Δ9 is even more surprising since Δ9 grows slightly slower than wild-type (1.36 ± 0.01 versus 1.46 ± 0.02 h^−1^ in rich medium for Δ9 and the wild-type strain, respectively)^2^. Hence, the cryptic prophages prevent premature persister resuscitation.

## DISCUSSION

As interwoven means of cell regulation, our results show that cryptic prophages not only help the cell respond to stress as an active response^2^ but also regulate the exit of the cell from dormancy. Specifically, we have demonstrated here that cryptic prophages control persister cell resuscitation through their inhibition of phosphate sensing. Clearly, although we found each of the nine cryptic prophages consistently inhibit persister resuscitation, cryptic prophage CP4-57 has the greatest effect (**Fig. 1B**), and regulator AlpA of CP4-57 was shown here to inhibit persister resuscitation. Since we discovered a 44-fold increase in waking with Δ9 and only a 4.4-fold increase with Δ*alpA*, there must be additional cryptic prophage proteins used by the host to reduce persister resuscitation. This is reasonable given that each of the nine cryptic prophages impacted resuscitation.

The mechanism for AlpA reducing resuscitation is likely by AlpA repressing *phoR*; *phoR* encodes the membrane-bound, sensor histidine kinase regulator of the phosphate-dependent PhoR/PhoB two-component signal transduction system that senses external phosphate (**Fig. 2**); PhoR/PhoB are active when concentrations of external phosphate are low^38^ by PhoR phosphorylating PhoB which induces the *pstSCAB* operon and facilitates phosphate uptake^39^. AlpA is a poorly-characterized, small (70 aa) regulator that likely binds DNA due its helix-turn-helix DNA-binding motif^40^; note the primary sequence only contains the DNA-binding motif due to its small size. AlpA is active in *E. coli* since it has been linked to AI-2 quorum-sensing in this strain (*alpA* is induced upon inactivation the AI-2 exporter TqsA^41^), and *alpA* is induced in *E. coli* mature biofilms^42^. Moreover, AlpA is a known transcriptional regulator since it causes excision of CP4-57^43^ by inducing *intA*^44^. Therefore, in the absence of AlpA, due to deleting cryptic prophage CP-457, derepression of *phoR* occurs, and we hypothesize that PhoR stimulates PhoB through phosphorylation, and phosphorylated PhoB increases persister cell resuscitation by reducing cAMP by silencing *cra* transcription^45, 46^, since Cra increases cAMP^47^. We found previously that decreasing cAMP dramatically increases resuscitation^3^ (**Fig. 2**).

Phosphate has been linked previously to persister cells formation, but, phosphate has not been studied previously for its impact on persister cell resuscitation. For *E. coli* persister cell formation, the phosphate regulator PhoU has been identified as a positive effector for persister cell formation^48^. However, the mechanism by which PhoU controls *E. coli* persister cell formation was not determined^49^, although in *S. aureus, phoU* deletion appears to reduce persistence by increasing slightly carbon metabolism^50^. PhoU regulates phosphate transport by repressing the phosphate-specific transport (Pst) system at high phosphate concentrations^49^. In addition, the *Mycobacterium tuberculosis* PhoU orthologs PhoY1 and PhoY2 also increase persister cell formation^49^. Therefore, our results on the cryptic prophages of *E. coli* reducing persister cell resuscitation fit well with these previous persister cell formation results related to phosphate since PhoU negatively regulates *phoR/phoB*^38^ (**Fig. 2**); i.e., PhoU increases persister cell *formation*^48, 49^, *and we found PhoB stimulates persister cell resuscitation*. Moreover, our results suggest a mechanism for the previous results indicating the negative regulator PhoU increases persistence: PhoU likely increases persistence by increasing cAMP through its inhibition of PhoB (**Fig. 2**).

The physiological benefit of reducing persister resuscitation appears to be to allow the cell to monitor more than phosphate concentrations before committing resources to waking. By deleting the cryptic prophages and negative regulators like AlpA, this highly-regulated return to active metabolism is short-circuited, leading to the dramatic increase seen in resuscitation, even in the absence of nutrients. Clearly, with limited ATP, the cell must optimize resuscitation (e.g., unlike exponentially-growing cells, persisters cannot wake without a carbon source in the presence of the cyptic prophages^3, 19^); hence, our results suggest the cell monitors the availability of carbon along with phosphate, before committing to resuscitation. Moreover, our results add credence to the idea that persister resuscitation is elegantly regulated^3^.

## MATERIALS AND METHODS

### Bacteria and growth conditions

Bacteria (**Table S5**) were cultured routinely in lysogeny broth^51^ at 37°C. The Δ9 strain lacking all nine *E. coli* cryptic prophage genes^2^ was verified through DNA microarrays to confirm that there were no undesired deletions and that each of the nine cryptic prophages was deleted completely^2^. The single cryptic prophage strains were verified by DNA sequencing^2^. The pCA24N-based plasmids^52^ were retained in overnight cultures via chloramphenicol (30 μg/mL), and kanamycin (50 μg/mL) was used for deletion mutants, where applicable. M9 glucose (0.4 wt%) medium^53^ (M9-Glu) was used for persister cell resuscitation.

### Persister cells

Exponentially-growing cells (turbidity of 0.8 at 600 nm) were converted to persister cells^9, 19^ by adding rifampicin (100 µg/mL) for 30 min to stop transcription, centrifuging, and adding LB with ampicillin (100 µg/mL) for 3 h to lyse non-persister cells. To remove ampicillin, cells were washed twice with 0.85% NaCl then re-suspended in 0.85% NaCl. Persister concentrations were enumerated via a drop assay^54^.

### ATP assay

To measure ATP concentrations in persister cells and in resuscitating cells, persister cells were formed with rifampicin/ampicillin treatment as indicated above, and for resuscitating cells, persister cells were resuspended in M9-Glu medium for 10 min. Samples (1 mL) were washed once and resuspended in Tris-acetate buffer (50 mM, pH 7.75), then the ATP assay was performed in duplicate using the ENLTEN ATP Assay System (Promega, cat#: FF2000) with the luminescence measured via a Turner Design 20E Luminometer using a 5 sec time delay and a 10 sec integration time.

### Single-cell persister resuscitation

Persister cells (5 µL) were added to 1.5% agarose gel pads containing M9-Glu medium, and single-cell resuscitation was visualized at 37°C via a light microscope (Zeiss Axio Scope.A1, bl_ph channel at 1000 ms exposure). For each condition, at least two independent cultures were used with 150 to 300 individual cells used per culture.

### RNA-Seq

To elucidate the transcriptome differences upon resuscitation of the Δ9 strain vs. wild-type, both strains were grown to a turbidity of 0.8 at 600 nm and converted into populations that solely consist of persisters by rifampicin and ampicillin treatment as indicated above, and resuscitated for 10 min in M9-Glu medium. Samples for RNA were added to cold 1.9 mL tubes containing RNA-Later, quick-cooled in dry ice-95% ethanol, centrifuged, and the cell pellets frozen at −80°C. RNA was harvested using the High Pure RNA Isolation Kit (Roche, Basel, Switzerland). Two independent samples were analyzed. The resultant library of RNA samples was quantified by a Bioanalyzer (Agilant) and sequenced by Illumina HiSeq 4000. Low quality reads and adapter sequences were filtered by cutadapt v2.8 [quality-cutoff (20), minimum-length (50)]^55^. Filtered reads were mapped to the reference genome (NZ_CP009273.1) using STAR v2.7.1a followed by ENCODE standard parameters^56^. Using the read mapping information obtained using the aligner, the expression levels of genes and transcripts were calculated using featureCounts v2.0.0, Cufflinks v2.2.1 (multi-read-correct, frag-bias-correct)^57, 58^, Differential expressed genes (DEG) were calculated by DESeq2 v1.26.0^59^

### qRT-PCR

To investigate the impact of AlpA production on phosphate sensing, qRT-PCR was performed for the phosphate sensing regulators *phoR, phoB* and *pstB* by producing AlpA from pCA24N-*alpA* in host Δ9 (compared to *Δ*9/pCA24N) that lacks AlpA in the chromosome. Cells were grown in LB until a turbidity of 0.4 at 600 nm, then *alpA* was induced with isopropyl β*-D*-1-thiogalactopyranoside (1 mM) for 90 min. RNA was isolated as for RNA-seq (above), *purM* was used as the housekeeping gene, and qRT-PCR was performed using Power SYBR Green RNA-to-CT 1-Step with a CFX96 Real-Time System and 100 ng of RNA along with the primers shown in **Table S6**.

## ACKNOWLEDGEMENTS

This work was supported by funds derived from the Biotechnology Endowed Professorship at the Pennsylvania State University for TKW, from the National Research Foundation of Korea (NRF) grant from the Korean Government (NRF-2020R1F1A1072397) for SYS, and from the National Research Foundation of Korea (NRF) funded by the Korean government (MSIT) (NRF-2019R1C1C1008856) for JSK. The authors have no conflicts of interest. Figure 2 was created with BioRender.com.

## Supporting Information

**Table S1.** ATP levels in persister cells and in resuscitating cells. Persister cells were formed by rifampicin pretreatment followed by ampicillin treatment for 3 h. For resuscitation, the persister cells were resuspended in M9-Glu medium for 10 min, then the ATP assay was performed. RLU is relative light units.

| **Persister cells** | **BW25113** | **$\Delta 9$** |
| --- | --- | --- |
| Average RLU/OD<sub>600</sub> | 15,000 ± 1700 | 11,000 ± 1600 |
| Relative RLU/OD<sub>600</sub> | 100% | -30% |
| Fold change | - | 0.7 |
| **Resuscitating cells** | **BW25113** | **$\Delta 9$** |
| Average RLU/OD<sub>600</sub> | 13,500 ± 500 | 6100 ± 200 |
| Relative RLU/OD<sub>600</sub> | 100% | -55% |
| Fold change | - | 0.45 |
| Relative to persisters, RLU/OD<sub>600</sub> | -11% | -43% |

**Table S2.** Single persister cell resuscitation on glucose gel pads. Single persister cells were observed using light microscopy (Zeiss Axio Scope.A1). The total number and waking number of persister cells are shown. Fold-change in resuscitation is relative to BW25113. These results are the combined observations from two independent experiments after 4 hours on 0.4 wt% glucose (independent culture results separated by “/”), and standard deviations are shown. The microscope images are shown in **Figure S1** and **Figure S3**.

|  | **Total**<br>**cells** | **Waking**<br>**cells** | **%**<br>**waking** | **Fold-**<br>**change** |
| --- | --- | --- | --- | --- |
| **BW25113** | 148/83 | 4/1 | 3 ± 5 | 1 |
| **Δ9** | 176/90 | 150/64 | 78 ± 10 | 40 |
| **Δ9**<br>(generated by treating<br>only with ampicillin) | 118/53 | 70/39 | 66 ± 10 | 34 |
| **Δe14** | 420/364 | 116/85 | 25 ± 3 | 13 |
| **ΔDLP-12** | 290/221 | 12/11 | 4.6 ± 0.6 | 2.3 |
| **ΔQin** | 293/122 | 55/33 | 23 ± 6 | 12 |
| **ΔCP4-6** | 190/241 | 33/44 | 17.8 ± 0.6 | 9 |
| **ΔRac** | 207/185 | 74/31 | 26 ± 13 | 14 |
| **ΔCPZ-55** | 278/199 | 69/35 | 21 ± 5 | 11 |
| **ΔCP4-44** | 163/58 | 15/18 | 20 ± 15 | 10 |
| **ΔCPS-53** | 170/99 | 19/0 | 6 ± 8 | 3 |
| **ΔCP4-57** | 284/133 | 103/79 | 48 ± 16 | 25 |
| **MG1655** | 382/237 | 11/5 | 2.5 ± 0.5 | 1 |
| **MG1655 MDS66** | 207/109 | 38/61 | 37 ± 27 | 17 |

**Table S3.** Single persister cell resuscitation on glucose gel pads. Single persister cells were observed using light microscopy (Zeiss Axio Scope.A1). The total number and waking number of persister cells are shown. Fold-changes in resuscitation are relative to BW25113/pCA24N lacking a cloned gene (Empty), relative to BW25113 (for Δ*alpA*), relative to Δ9/pCA24N (for Δ9/AlpA), or relative to BW25113 on gel pads that lack nutrients (no nutrients). These results are the combined observations from two independent experiments for 6 h for BW25113/pCA24N strains, for 4 h for BW25113 vs. Δ*alpA*, and for 3 h for Δ9/pCA24N strains and BW25113 (no nutrients) vs. Δ9 (no nutrients); the different resuscitation times were used to distinguish more clearly the differences in resuscitation. Resuscitation was at 37°C on 0.4 wt% glucose except for the “no nutrients” experiments where glucose was not used. Independent culture results are separated by “/”, and standard deviations are shown. The microscope images are shown in **Figure S4**.

|  | Total<br>cells | Waking<br>cells | % waking | Fold-change |
| --- | --- | --- | --- | --- |
| **Empty** | 205/119 | 5/2 | 2.1 ± 0.5 | 1 |
| **PsiF** | 130/36 | 13/12 | 22 ± 17 | 4.8 |
| **PhoB** | 243/153 | 35/20 | 14 ± 1 | 7.7 |
| **PhoU** | 270/83 | 2/0 | 0.4 ± 0.5 | -4.0 |
| **$\Delta alpA$** | 755/539 | 214/98 | 23 ± 7 | 4.4 |
| **$\Delta 9$/Empty** | 559/476 | 393/185 | 54.6 ± 22.2 | 1 |
| **$\Delta 9$/AlpA** | 276/221 | 62/17 | 14.9 ± 10.2 | -4.1 |
| **BW25113 (no nutrients)** | 278/271 | 0/0 | 0 | 1 |
| **$\Delta 9$ (no nutrients)** | 235/211 | 19/22 | 9.3 ± 2.0 | ∞ |

**Table S4.** Repression of *pstB, phoB*, and *phoR* by AlpA in the Δ9 host. Fold changes (normalized by housekeeping gene *purM*) were calculated relative to the transcription level detected for the empty plasmid pCA24N in host Δ9. Cells were grown in LB until a turbidity of 0.4 at 600 nm, then *alpA* was induced with isopropyl β*-D*-1-thiogalactopyranoside (1 mM) for 90 min.

| Gene | Strain | $\Delta C_T$ | $\Delta \Delta C_T$ | fold-change |
| --- | --- | --- | --- | --- |
| *pstB* | $\Delta 9$/pCA24N-*alpA* | $1.4 \pm 4$ | $3.0 \pm 4$ | -8 |
| | $\Delta 9$/pCA24N | $-1.5 \pm 2$ | | |
| *phoB* | $\Delta 9$/pCA24N-*alpA* | $6.9 \pm 3$ | $1.2 \pm 3$ | -2.3 |
| | $\Delta 9$/pCA24N | $5.7 \pm 1$ | | |
| *phoR* | $\Delta 9$/pCA24N-*alpA* | $3.4 \pm 3$ | $2.9 \pm 3$ | -7.3 |
| | $\Delta 9$/pCA24N | $0.57 \pm 1$ | | |

**Table S5.**
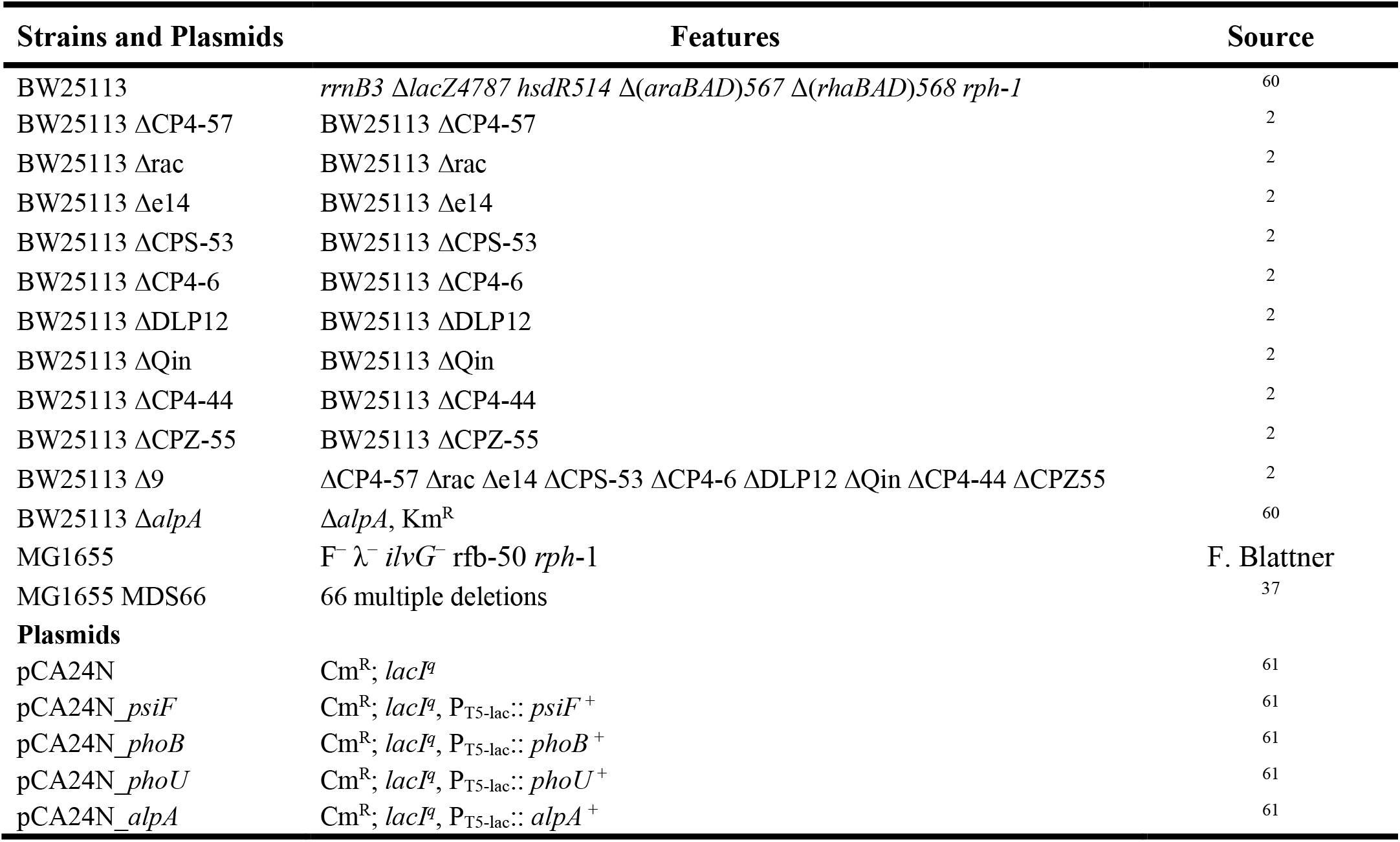
*E. coli* bacterial strains. Bacterial strains and plasmids used in this study.

**Table S6.**
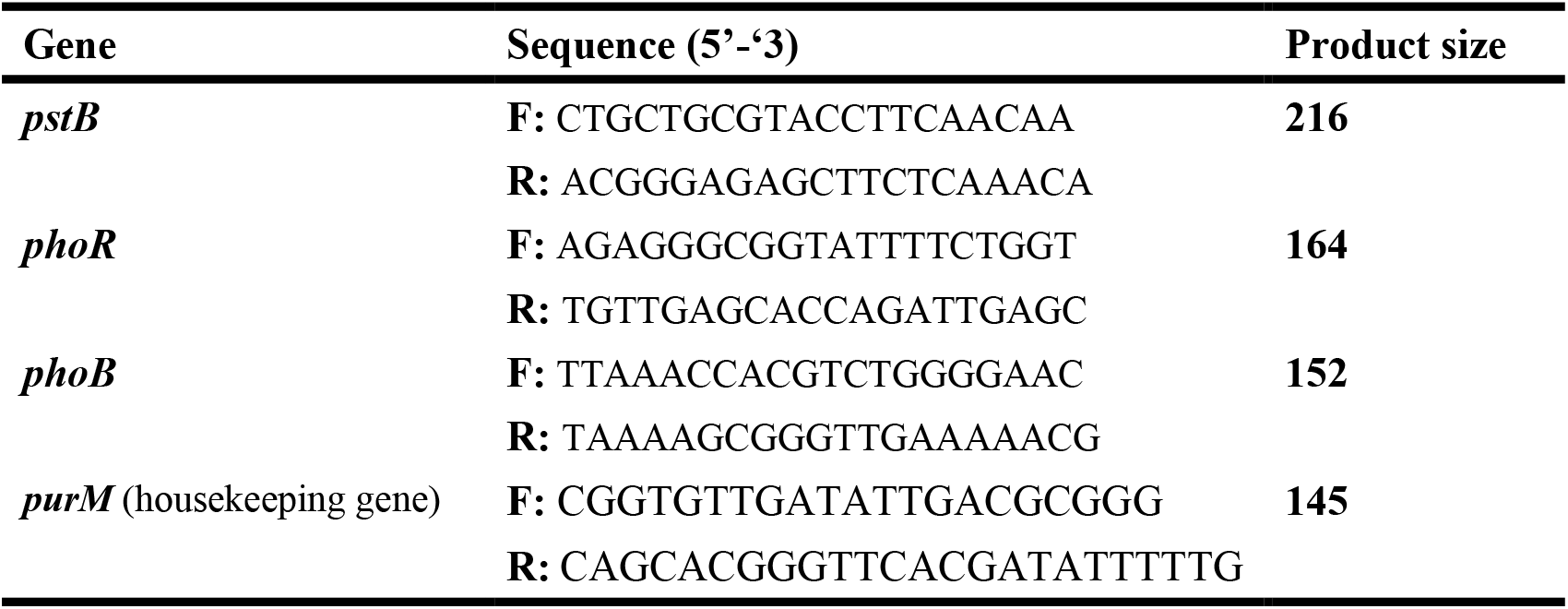
Primer sequences used for qRT-PCR.

**Supplementary Figure 1.**
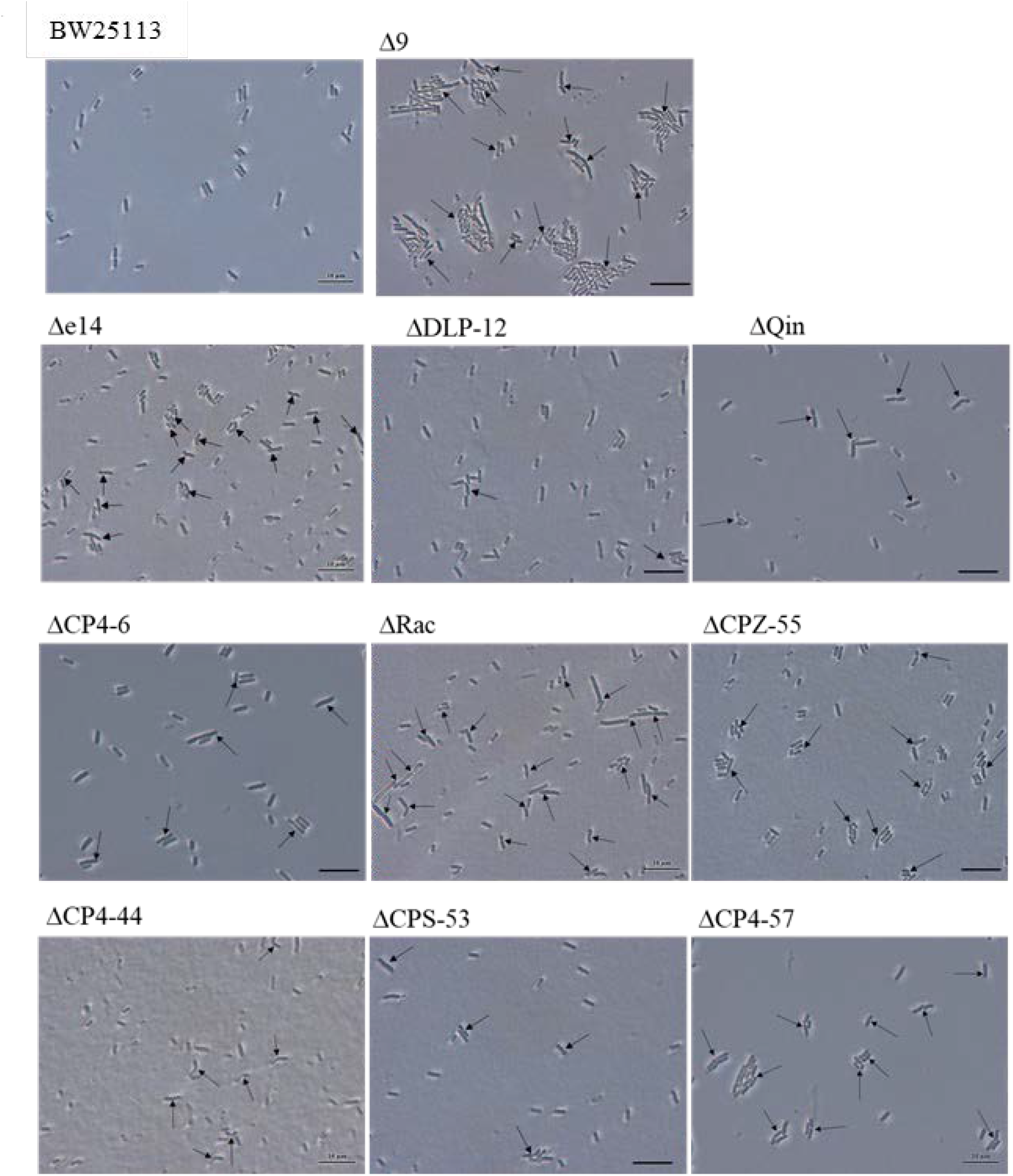
Single cell persister waking of BW25113, BW25113 Δ9, and single cryptic prophage deletions BW25113 Δe14, BW25113 ΔDLP-12, BW25113 ΔQin, BW25113 ΔCP4-6, BW25113 ΔRac, BW25113 ΔCPZ-55, BW25113 ΔCP4-44, BW25113 ΔCPS-53, and BW25113 ΔCP4-57. Single-cell persister resuscitation after 4 h at 37°C with 0.4 wt% glucose compared to the wild-type strain as determined using light microscopy (Zeiss Axio Scope.A1). Black arrows indicate cells that resuscitate. Scale bar indicates 10 µm. Representative results from two independent cultures are shown, and tabulated cell numbers are shown in **Table S2**.

**Supplementary Figure 2.**
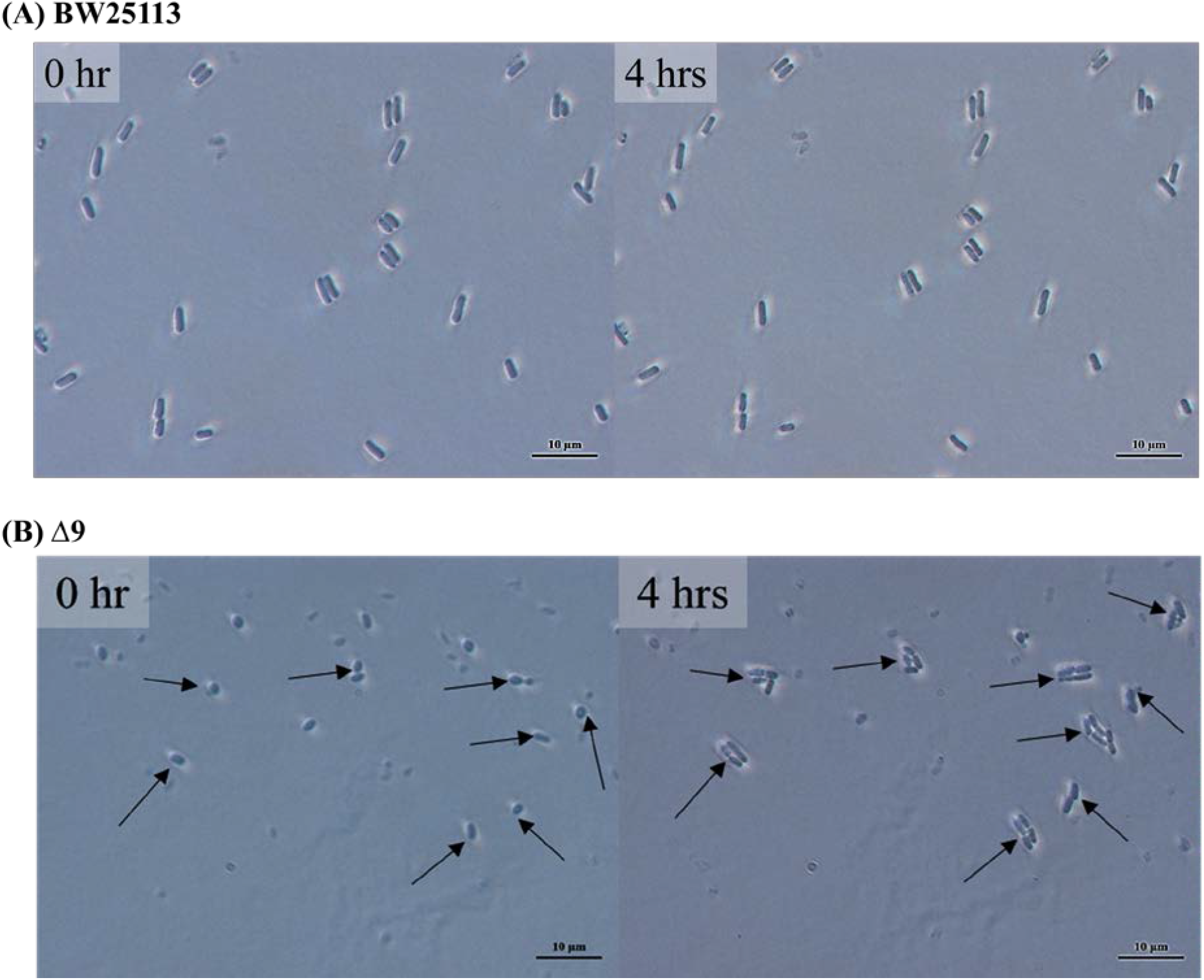
Single cell persister waking of BW25113 Δ9 with the persister state induced only with ampicillin. Persister cell waking of BW25113 (**A**) and BW25113 Δ9 (**B**) on M9 0.4% glucose agar plates incubated at 37°C for 4 hours as determined using light microscopy (Zeiss Axio Scope.A1). Black arrows indicate all waking cells. Scale bars indicate 10 µm. Representative results from two independent cultures are shown and tabulated cell numbers are shown in **Table S2**. Persister cells were generated by treating only with ampicillin for 3 h (no rifampicin pretreatment).

**Supplementary Figure 3.**
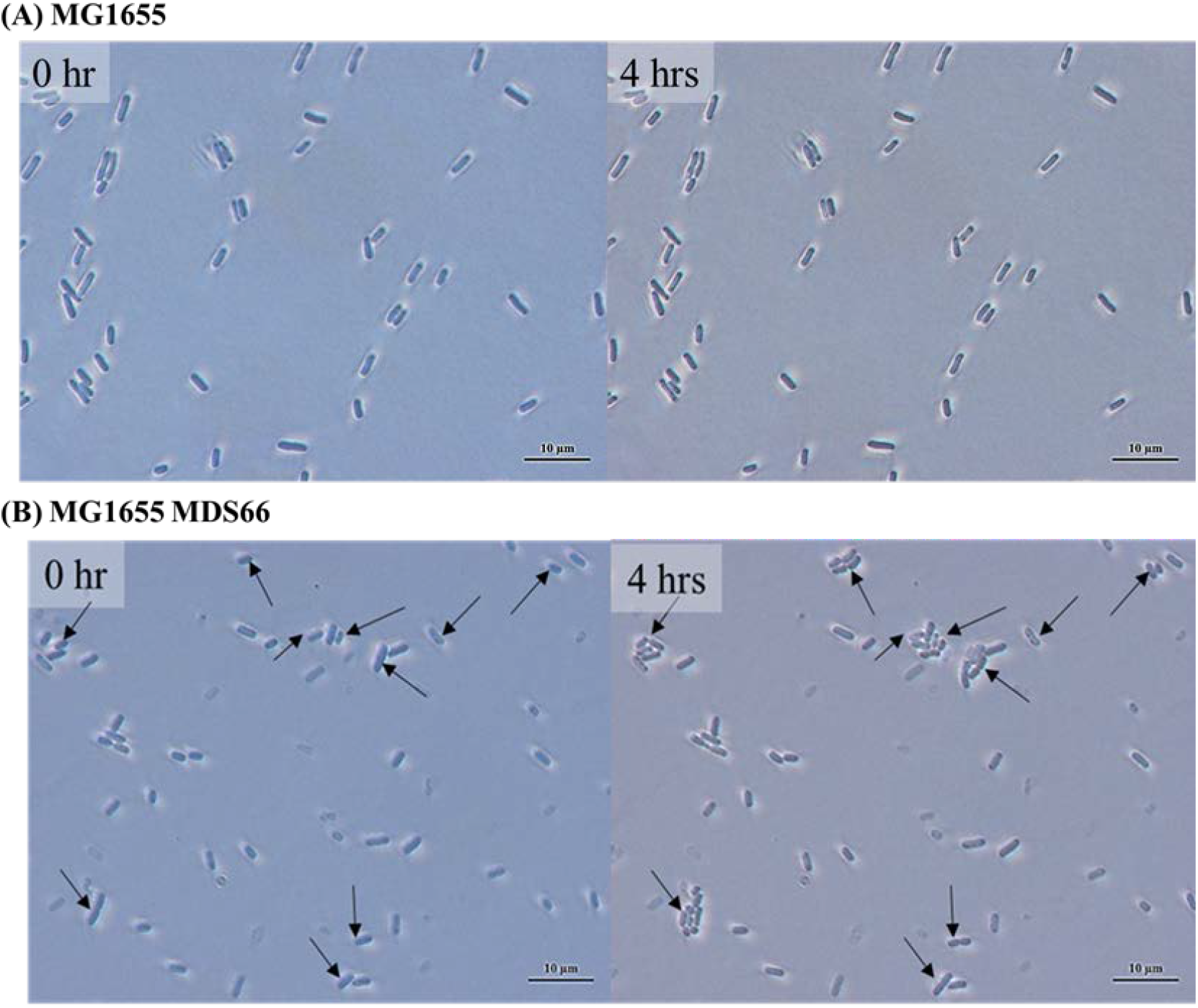
Single cell persister waking of deletion strain MG1655 MDS66. Persister cell waking of MG1655 (**A**) and MG1655 MDS66 (**B**) on M9 0.4% glucose agar plates incubated at 37°C for 4 hours as determined using light microscopy (Zeiss Axio Scope.A1). Black arrows indicate all waking cells. Scale bars indicate 10 µm. Representative results from two independent cultures are shown and tabulated cell numbers are shown in **Table S2**.

**Supplementary Figure 4.**
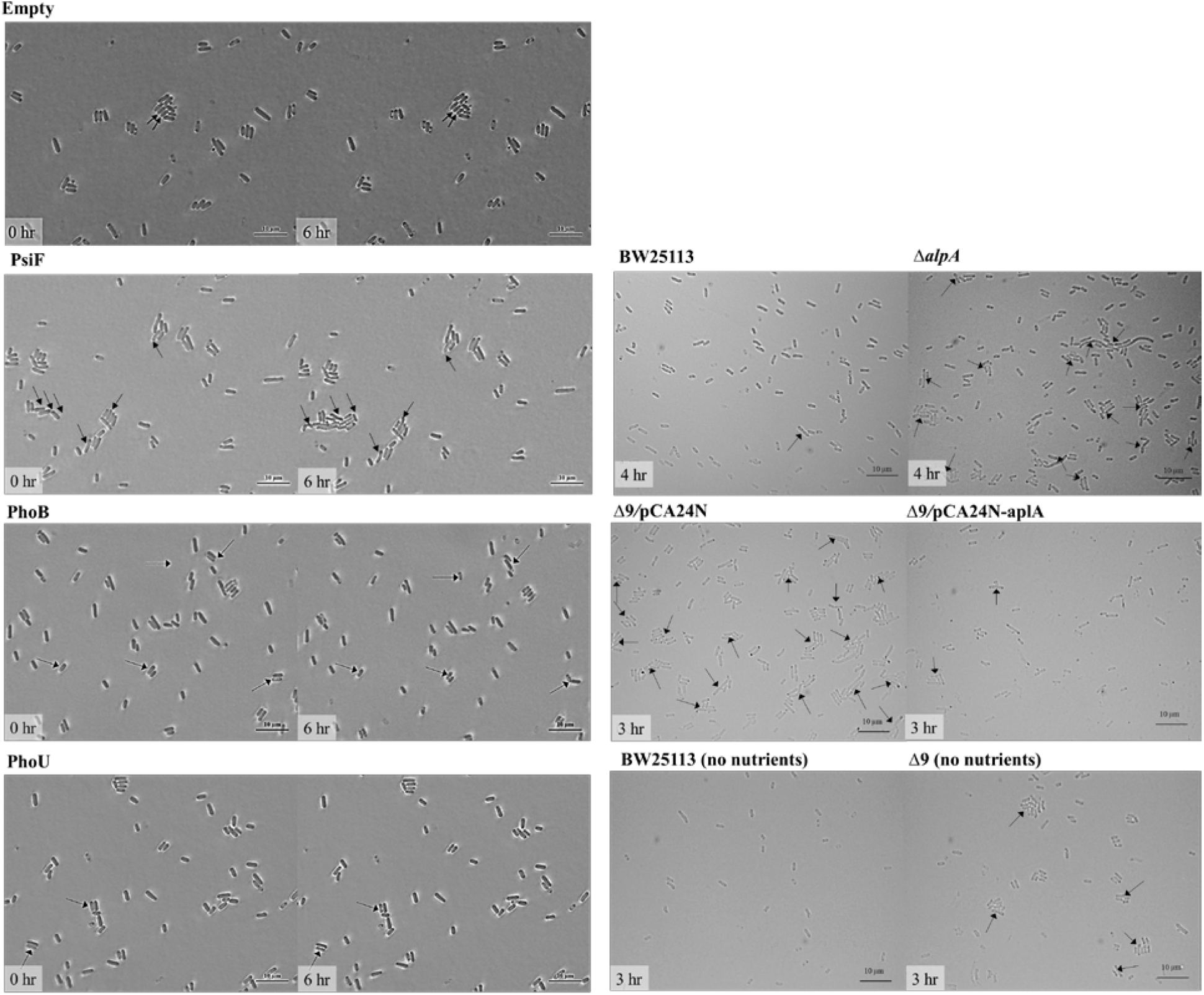
Single-cell persister waking after producing PhoB and PhoU, after deleting *alpA*, after producing AlpA, and after removing glucose (no nutrients). Persister cell waking in 6 h for BW25113/pCA24N (“Empty”) vs. BW25113/pCA24N_*psiF* (“PsiF”), BW25113/pCA24N (“Empty”) vs. BW25113/pCA24N_*phoB* (“PhoB”), and BW25113/pCA24N (“Empty”) vs. BW25113/pCA24N_*phoU* (“PhoU”), after 4 h for BW25113 vs. BW25113 Δ*alpA*, after 3 h for Δ9/pCA24N vs. Δ9/pCA24N_*alpA*, and after 3 h for BW25113 (no nutrients) vs. BW25113 Δ9 (no nutrients); the different resuscitation times were used to distinguish more clearly the differences in resuscitation. Cells were resuscitated at 37°C with 0.4 wt% glucose (except for the “no nutrient” group) as determined using light microscopy (Zeiss Axio Scope.A1). Black arrows indicate cells that resuscitate. Scale bar indicates 10 µm. Representative results from two independent cultures are shown, and tabulated cell numbers are shown in **Table S3**.

## REFERENCES

1. Casjens, S. Prophages and bacterial genomics: what have we learned so far? Mol. Microbiol. 49, 277–300 (2003).

2. Wang, X. et al. Cryptic prophages help bacteria cope with adverse environments. Nat Commun 1, 147 (2010).

3. Yamasaki, R., Song, S., Benedik, M.J. & Wood, T.K. Persister Cells Resuscitate Using Membrane Sensors that Activate Chemotaxis, Lower cAMP Levels, and Revive Ribosomes. iScience 23, 100792 (2020).

4. Hong, S.H., Wang, X., O’Connor, H.F., Benedik, M.J. & Wood, T.K. Bacterial persistence increases as environmental fitness decreases. Microbial Biotechnol 5, 509–522 (2012).

5. Kim, J.-S., Chowdhury, N., Yamasaki, R. & Wood, T.K. Viable But Non-Culturable and Persistence Describe the Same Bacterial Stress State. Environ Microbiol 20, 2038–2048 (2018).

6. Lewis, K. Multidrug tolerance of biofilms and persister cells. Curr Top Microbiol Immunol 322, 107–131 (2008).

7. Song, S. & Wood, T.K. Post-segregational Killing and Phage Inhibition Are Not Mediated by Cell Death Through Toxin/Antitoxin Systems. Front Microbiol 9, 814 (2018).

8. Van den Bergh, B., Fauvart, M. & Michiels, J. Formation, physiology, ecology, evolution and clinical importance of bacterial persisters. FEMS MIcrobiol Rev 41, 219–251 (2017).

9. Kwan, B.W., Valenta, J.A., Benedik, M.J. & Wood, T.K. Arrested protein synthesis increases persister-like cell formation. Antimicrob Agents Chemother 57, 1468–1473 (2013).

10. Korch, S.B., Henderson, T.A. & Hill, T.M. Characterization of the hipA7 allele of Escherichia coli and evidence that high persistence is governed by (p)ppGpp synthesis. Mol Microbiol 50, 1199–1213 (2003).

11. Chowdhury, N., Kwan, B.W. & Wood, T.K. Persistence Increases in the Absence of the Alarmone Guanosine Tetraphosphate by Reducing Cell Growth. Sci Rep 6, 20519 (2016).

12. Nguyen, D. et al. Active starvation responses mediate antibiotic tolerance in biofilms and nutrient-limited bacteria. Science 334, 982–986 (2011).

13. Svenningsen, M.S., Veress, A., Harms, A., Mitarai, N. & Semsey, S. Birth and Resuscitation of (p)ppGpp Induced Antibiotic Tolerant Persister Cells. Sci Rep 9, 6056 (2019).

14. Shimada, T., Yoshida, H. & Ishihama, A. Involvement of Cyclic AMP Receptor Protein in Regulation of the rmf Gene Encoding the Ribosome Modulation Factor in Escherichia coli. J Bacteriol 195, 2212–2219 (2013).

15. Song, S. & Wood, T.K. ppGpp Ribosome Dimerization Model for Bacterial Persister Formation and Resuscitation. Biochem. Biophys. Res. Com. 523, 281–286 (2020).

16. Wood, T.K. & Song, S. Forming and waking dormant cells: The ppGpp ribosome dimerization persister model. Biofilm 2, 100018 (2020).

17. Song, S. & Wood, T.K. Persister Cells Resuscitate via Ribosome Modification by 23S rRNA Pseudouridine Synthase RluD. Environ. Microbiol. 22, 850–857 (2020).

18. Goormaghtigh, F. et al. Reassessing the Role of Type II Toxin-Antitoxin Systems in Formation of Escherichia coli Type II Persister Cells. mBio 9, e00640–00618 (2018).

19. Kim, J.-S., Yamasaki, R., Song, S., Zhang, W. & Wood, T.K. Single Cell Observations Show Persister Cells Wake Based on Ribosome Content. Environ Microbiol 20, 2085–2098 (2018).

20. Pu, Y. et al. ATP-Dependent Dynamic Protein Aggregation Regulates Bacterial Dormancy Depth Critical for Antibiotic Tolerance. Molecular Cell 73, 143–156 (2019).

21. Goormaghtigh, F. & Van Melderen, L. Single-cell imaging and characterization of Escherichia coli persister cells to ofloxacin in exponential cultures. Science Advances 5, eaav9462 (2019).

22. Mohiuddin, S.G., Kavousi, P. & Orman, M.A. Flow-cytometry analysis reveals persister resuscitation characteristics. BMC Microbiology 20, 202 (2020).

23. Cui, P. et al. Identification of Genes Involved in Bacteriostatic Antibiotic-Induced Persister Formation. Front Microbiol 9, 413 (2018).

24. Grassi, L. et al. Generation of Persister Cells of Pseudomonas aeruginosa and Staphylococcus aureus by Chemical Treatment and Evaluation of Their Susceptibility to Membrane-Targeting Agents. Front Microbiol 8, 1917 (2017).

25. Narayanaswamy, V.P. et al. Novel Glycopolymer Eradicates Antibiotic-and CCCP-Induced Persister Cells in Pseudomonas aeruginosa. Front Microbiol 9, 1724 (2018).

26. Sulaiman, J.E., Hao, C. & Lam, H. Specific Enrichment and Proteomics Analysis of Escherichia coli Persisters from Rifampin Pretreatment. J Proteome Res 17, 3984–3996 (2018).

27. Tkhilaishvili, T., Lombardi, L., Klatt, A.-B., Trampuz, A. & Di Luca, M. Bacteriophage Sb-1 enhances antibiotic activity against biofilm, degrades exopolysaccharide matrix and targets persisters of Staphylococcus aureus. Int J Antimicrob Agents 52, 842–853 (2018).

28. Rowe, S.E. et al. Reactive oxygen species induce antibiotic tolerance during systemic Staphylococcus aureus infection. Nature Microbiology 5, 282–290 (2020).

29. Zhao, Y. et al. Rapid Freezing Enables Aminoglycosides To Eradicate Bacterial Persisters via Enhancing Mechanosensitive Channel MscL-Mediated Antibiotic Uptake. mBio 11, e03239–03219 (2020).

30. Yu, W. et al. Absence of tmRNA Increases the Persistence to Cefotaxime and the Intercellular Accumulation of Metabolite GlcNAc in Aeromonas veronii. Frontiers in Cellular and Infection Microbiology 10 (2020).

31. Sun, F. et al. 5-Methylindole Potentiates Aminoglycoside Against Gram-Positive Bacteria Including Staphylococcus aureus Persisters Under Hypoionic Conditions. Frontiers in Cellular and Infection Microbiology 10, 84 (2020).

32. Zheng, E.J., Stokes, J.M. & Collins, J.J. Eradicating Bacterial Persisters with Combinations of Strongly and Weakly Metabolism-Dependent Antibiotics. Cell Chemical Biology 27, P1544-1552.E1543 (2020).

33. Jin, X., Kightlinger, W., Kwon, Y.-C. & Hong, S.H. Rapid production and characterization of antimicrobial colicins using Escherichia coli-based cell-free protein synthesis. Synthetic Biology 3, ysy004 (2018).

34. Martins, D., McKay, G.A., English, A.M. & Nguyen, D. Sublethal Paraquat Confers Multidrug Tolerance in Pseudomonas aeruginosa by Inducing Superoxide Dismutase Activity and Lowering Envelope Permeability. Front. Microbiol. 11, 576708 (2020).

35. Johnson, P.J.T. & Levin, B.R. Pharmacodynamics, Population Dynamics, and the Evolution of Persistence in Staphylococcus aureus. PLOS Genetics 9, e1003123 (2013).

36. Harms, A., Fino, C., Sørensen, M.A., Semsey, S. & Gerdes, K. Prophages and Growth Dynamics Confound Experimental Results with Antibiotic-Tolerant Persister Cells. mBio 8, e01964–01917 (2017).

37. Karcagi, I. et al. Indispensability of Horizontally Transferred Genes and Its Impact on Bacterial Genome Streamlining. Molecular Biology and Evolution 33, 1257–1269 (2016).

38. Kritmetapak, K. & Kumar, R. Phosphate as a Signaling Molecule. Calcified Tissue International (2019).

39. Santos-Beneit, F. The Pho regulon: a huge regulatory network in bacteria. Front. Microbiol. 6, 402 (2015).

40. Keseler, I.M. et al. The EcoCyc database: reflecting new knowledge about Escherichia coli K-12. Nucleic Acids Research 45, D543–D550 (2016).

41. Herzberg, M., Kaye, I.K., Peti, W. & Wood, T.K. YdgG (TqsA) Controls Biofilm Formation in Escherichia coli K-12 by Enhancing Autoinducer 2 Transport. J. Bacteriol. 188, 587–598 (2006).

42. Domka, J., Lee, J., Bansal, T. & Wood, T.K. Temporal gene-expression in Escherichia coli K-12 biofilms. Environ. Microbiol. 9, 332–346 (2007).

43. Wang, X., Kim, Y. & Wood, T.K. Control and Benefits of CP4-57 Prophage Excision in Escherichia coli Biofilms. ISME J 3, 1164–1179 (2009).

44. Kirby, J.E., Trempy, J.E. & Gottesman, S. Excision of a P4-like cryptic prophage leads to Alp protease expression in Escherichia coli. Journal of Bacteriology 176, 2068–2081 (1994).

45. Marzan, L.W. & Shimizu, K. Metabolic regulation of Escherichia coli and its phoB and phoR genes knockout mutants under phosphate and nitrogen limitations as well as at acidic condition. Microbial Cell Factories 10, 39 (2011).

46. Marzan, L.W., Hasan, C.M.M. & Shimizu, K. Effect of acidic condition on the metabolic regulation of Escherichia coli and its phoB mutant. Archives of Microbiology 195, 161–171 (2013).

47. Crasnier-Mednansky, M., Park, M.C., Studley, W.K. & Saier, M.H. Cra-mediated regulation of Escherichia coli adenylate cyclase. Microbiology 143, 785–792 (1997).

48. Li, Y. & Zhang, Y. PhoU Is a Persistence Switch Involved in Persister Formation and Tolerance to Multiple Antibiotics and Stresses in Escherichia coli. Antimicrobial Agents and Chemotherapy 51, 2092–2099 (2007).

49. Namugenyi, S.B., Aagesen, A.M., Elliott, S.R. & Tischler, A.D. Mycobacterium tuberculosis PhoY Proteins Promote Persister Formation by Mediating Pst/SenX3-RegX3 Phosphate Sensing. mBio 8, e00494–00417 (2017).

50. Shang, Y. et al. Staphylococcus aureus PhoU Homologs Regulate Persister Formation and Virulence. Front. Microbiol. 11, 865 (2020).

51. Bertani, G. Studies on Lysogenesis .1. The Mode of Phage Liberation by Lysogenic Escherichia-Coli. J Bacteriol 62, 293–300 (1951).

52. Kitagawa, M. et al. Complete set of ORF clones of Escherichia coli ASKA library (A Complete Set of E. coli K-12 ORF Archive): Unique Resources for Biological Research. DNA Res 12, 291–299 (2005).

53. Rodriguez, R.L. & Tait, R.C. Recombinant DNA Techniques: An Introduction. (Benjamin/Cummings Publishing, Menlo Park, CA; 1983).

54. Donegan, K., Matyac, C., Seidler, R. & Porteous, A. Evaluation of methods for sampling, recovery, and enumeration of bacteria applied to the phylloplane. Appl Environ Microbiol 57, 51–56 (1991).

55. Martin, M. Cutadapt removes adapter sequences from high-throughput sequencing reads. EMBnet.journal 17, 1 (2011).

56. Dobin, A. et al. STAR: ultrafast universal RNA-seq aligner. Bioinformatics 29, 15–21 (2012).

57. Liao, Y., Smyth, G.K. & Shi, W. featureCounts: an efficient general purpose program for assigning sequence reads to genomic features. Bioinformatics 30, 923–930 (2013).

58. Trapnell, C. et al. Transcript assembly and quantification by RNA-Seq reveals unannotated transcripts and isoform switching during cell differentiation. Nature Biotechnology 28, 511–515 (2010).

59. Love, M.I., Huber, W. & Anders, S. Moderated estimation of fold change and dispersion for RNA-seq data with DESeq2. Genome Biology 15, 550 (2014).

60. Baba, T. et al. Construction of Escherichia coli K-12 in-frame, single-gene knockout mutants: the Keio collection. Mol. Syst. Biol. 2, 2006 0008 (2006).

61. Kitagawa, M. et al. Complete set of ORF clones of Escherichia coli ASKA library (a complete set of E. coli K-12 ORF archive): unique resources for biological research. DNA Res. 12, 291–299 (2005).

